# H3K9 methyltransferase EHMT2/G9a controls ERVK-driven non-canonical imprinted genes

**DOI:** 10.1101/2021.03.29.437617

**Authors:** Tie-Bo Zeng, Nicholas Pierce, Piroska E. Szabó

## Abstract

Unlike regular imprinted genes, non-canonical imprinted genes are known to not depend on gamete-specific DNA methylation difference. Instead, the paternal allele-specific expression of these genes in the extra-embryonic lineages depends on an H3K27me3-based imprint in the oocyte, but this marking is not maintained beyond pre-implantation development. The maintenance of non-canonical imprinting corresponds to maternal allele-specific DNA methylation and paternal allele-specific H3K4me3 at their somatic DMRs, which occur at ERVK repeats. We hypothesized that EHMT2, the main euchromatic H3K9 methyltransferase, also has a role in this process. Using reciprocal mouse crosses and allele-specific RNA-seq analysis, we found that the maternal allele of each known non-canonical imprinted gene was derepressed from its ERVK promoter in the *Ehmt2*^*−/−*^ ectoplacental cone of somite-matched 8.5 dpc embryos. In the *Ehmt2*^*−/−*^ embryos, loss of DNA methylation accompanied the derepression of both parental alleles of those ERVK promoters. Our study identifies EHMT2 as an essential player that maintains the repressed chromosomal state in non-canonical imprinting.

## Introduction

Genomic imprinting is an epigenetic process that directs genes to be expressed in a parental allele-specific manner (*1, 2*). Imprinted genes frequently exist in clusters and respond to the domain-wide control by germline-derived differentially methylated regions (gDMRs) (*3*). The parental allele-specific differential DNA methylation at gDMRs is established during gametogenesis and is maintained in the zygote, where global DNA demethylation takes place, and continues to be maintained during preimplantation and throughout development. Mutations that disrupt the *de novo* DNA methylation process during male or female gametogenesis cause the loss of paternal or maternal methylation imprints at the gDMRs and the loss of parental allele-specific expression of imprinted genes in the soma (*4, 5*).

A set of imprinted genes is specifically imprinted in the placenta but not in the embryo (*6–8*). Several of these genes have been previously demonstrated to play important roles in placenta function and development (*9*). Their exact gene dosage is often critical, which is controlled by genomic imprinting in the placenta (*9*). Non-canonical imprinted genes are a special group of placenta-specific imprinted genes. The establishment of non-canonical imprinting is independent of oocyte-specific DNA methylation, as conditional knockout of *Dnmt3a/3b* or *Dnmt3l* in the growing oocytes did not affect the paternal allele-specific expression of these genes in the extra-embryonic lineage (*10, 11*). The key gametic imprinting mark of non-canonical imprinted genes in the oocyte is the repressive H3K27me3 modification at their promoters, which is required for imprinted expression of these genes in the trophectoderm of the blastocyst and later in the extraembryonic ectoderm (*12*).

Because the maternal allele-specific H3K27me3 is not maintained beyond pre-implantation development (*10*), the maintenance of non-canonical imprinting requires additional mechanisms. Indeed, such a maintenance role was shown for DNA methylation, when zygotic double knockout of *Dnmt3a/3b* led to loss of silencing of the maternal allele of non-canonical imprinted genes during post-implantation development (*11*). More recently, the activity of alternative promoters have been discovered at non-canonical imprinted genes, each coinciding with the long terminal repeat (LTR) of an endogenous retrovirus-K (ERVK) element (*10*). These ERVK promoters become de novo methylated in the maternal allele in the extra-embryonic ectoderm, and in both alleles in the epiblast at 6.5 days post coitum (dpc) (*10*). It is not known what controls the allele-specificity of DNA methylation at these secondary DMRs (sDMRs).

G9a/EHMT2 functions as the main H3K9 methyltransferase in euchromatin (*13*). It is essential for embryo development, as zygotic mutant *Ehmt2* null embryos die at around 10.5 dpc (*14, 15*) and conditional maternal knockout of *Ehmt2* in the growing oocytes causes severe developmental delay of preimplantation-stage embryos (*16, 17*). Because EHMT2 is known to direct DNA methylation to transposable elements (*2, 18–20*), and to play important roles in placenta development and maintaining canonical imprinting in the placenta (*21*), we hypothesized that it may have a role in regulating ERVKs and non-canonical imprinting. In this study we performed allele-specific total RNA-seq analysis of maternal and zygotic *Ehmt2*-mutant ectoplacental cones (EPCs), and embryos to test the role of EHMT2 in controlling non-canonical imprinted genes.

## Results

### EHMT2 is required for the maintenance of non-canonical imprinting in the placenta

We hypothesized that EHMT2 enzymatic activity is required for maintaining the paternal allele-specific expression of non-canonical imprinted genes. We made use of an *Ehmt2* mutant mouse line in which the exons encoding the SET domain were deleted (*22*). The mutation eliminated the catalytic activity of the EHMT2 protein, as judged by the significantly reduced intensity of H3K9me2 staining in the maternal pronucleus of the *Ehmt2*^m-z+^ zygotes (*22*). We decided to investigate noncanonical imprinting in the 8.5 dpc EPC. We carried out RNAseq experiments using the EPCs of *Ehmt2*^−/−^ homozygous (HOMO) and *Ehmt2*^*+/+*^ wild type (WT) embryos, where the parental genomes were distinguished by single nucleotide polymorphisms (SNPs) between the 129S1 (129) and JF1/Ms (JF1) inbred mouse strains (Fig. 1A). These samples were derived by crossing *Ehmt2*^+/−(JF1.N10)^ females, where the mutation has been backcrossed into the JF1 background, with *Ehmt2*^+/−(129)^ males (JF1 × 129), and by crossing the reciprocal parents (129 JF). In addition, we also generated *Ehmt2*^+/+^ (CONTROL) EPCs from *Ehmt2*^+/+^ parents (JF1×129 and 129xJF) to account for any potential effects that can arise from parental haploinsufficiency. We found that the majority of HOMO embryos reach the 6-somite stage at the time when WT embryos already reach the 12-somite stage around 8.5 days post coitum (dpc). We designate these HOMO6 and WT12, respectively. To control for the developmental delay due to the mutation, we included EPCs not only from age-matched uterus mate embryos (HOMO6 and WT12), but also from developmental stage-matched embryos (WT6 and HOMO12) collected at different times of gestation (Table S1). WT6 and HOMO6 samples were collected from reciprocal crosses and WT12 and HOMO12 samples were collected from the JF1×129 cross (Fig. 1A).

**Fig. 1.**
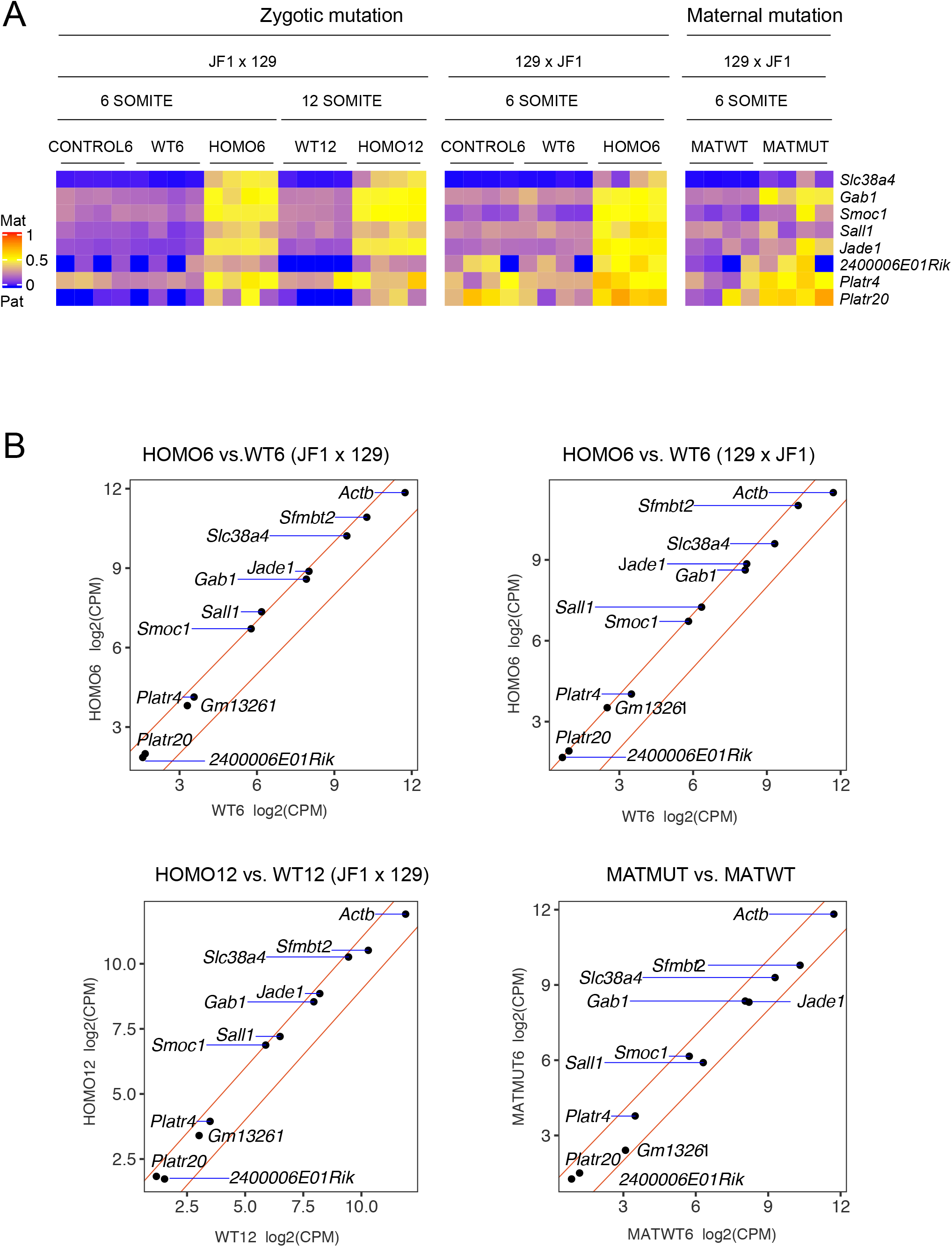
EHMT2 controls non-canonical imprinting in the ectoplacental cone. **(A)** Allelic expression ratios (Mat/Mat+Pat) of selected known and novel non-canonical imprinted genes are depicted across the EPC samples as marked above. The heatmap shows the results of allele-specific RNA-seq experiment using reciprocal 129×JF1 and JF1×129 (female x male) mouse crosses between mice of the 129S1 (129) and the JF1/Ms (JF1) inbred background to allow distinguishing the parental alleles by SNPs. EPCs were collected from *Ehmt2*^−/−^ zygotic mutant (HOMO), and *Ehmt2*^mat−/+^ maternal mutant (MATMUT) embryos, and their matching controls that reached the 6-somite or 12-somite stages, as indicated. In addition to the *Ehmt2*^+/+^ wild type (WT) embryos derived from two *Ehmt2*^+/−^ heterozygous parents we also generated *Ehmt2*^+/+^ (CONTROL) embryos derived from *Ehmt2*^+/+^ parents. Four replicates (two female and two male) were used for each genotype at each stage. The details of the samples can be found in Table S1. **(B)** Relative expression levels of non-canonical imprinted genes in the mutant versus wild-type EPCs are displayed by scatter plots as marked. The average values from four replicates were used for the differential gene expression analysis. The red lines mark the +2-fold and −2-fold difference cutoff values.

We performed an allele-specific and strand-specific total RNAseq experiment using four replicate EPCs, two females and two males, per condition (listed in Table S1). A gene was determined as maternally or paternally expressed using the cutoff allelic expression ratio [mat/mat+pat] of > 0.7 or < 0.3, respectively. We found that the known non-canonical imprinted genes *Slc38a4*, *Gab1*, *Smoc1*, *Sall1*, *and Jade1 (Phf17)* were paternally expressed in the CONTROL and WT EPCs in the JF1 × 129 and the reciprocal 129×JF1 cross at the 6-somite and also the 12-somite stages, as depicted in the heatmap in Figure 1A. The allelic expression bias of *Platr20* was stronger in the WT EPCs from the JF1×129 cross than from the reciprocal 129S x JF1 cross (Fig. 1A), indicating that some degree of strain-specific bias exists for this known non-canonical imprinted gene. In the absence of EHMT2, these genes were all biallelically expressed in the HOMO EPCs in the JF1×129 and the reciprocal 129×JF1 cross. These data show that zygotic *Ehmt2* mutation causes loss of imprinting of known non-canonical imprinted genes in the HOMO EPC and also the loss of paternal bias at *Platr20*. By calling imprinted genes that significantly change allelic expression ratio in the HOMO samples compared to the WT samples (FDR>0.05) (Table S2) we found an additional gene *Platr4* that behaved similarly to the known noncanonical imprinted genes. Consistent with the biallelic expression, the expression levels of the non-canonical imprinted genes were upregulated ~2-fold in the HOMO EPCs at the 6-somite stage in both JF1×129 and 129×JF1 crosses, and also at the 12-somite stage from JF1×129 cross (Fig. 1B). Collectively, these results demonstrate that the maintenance of paternal allele-specific expression of the non-canonical imprinted genes in the EPC requires EHMT2 catalytic activity.

### Maternal EHMT2 may be required for establishing non-canonical imprinting

We hypothesized that EHMT2 may also be required in the oocyte for establishing the non-methylation maternal imprinting at the non-canonical imprinted genes. To test this possibility, we analyzed *Ehmt2*^*mat*−/+^ maternal mutant (MATMUT) EPCs (Fig. 1A). The MATMUT EPCs were generated by crossing *Ehmt2*^*fl/fl*^ (129); *Zp3-Cre*^*Tg/0*^ females with wild type males (JF1). The *Zp3-cre* transgene-derived Cre recombinase excises the floxed SET domain-coding exons of *Ehmt2* in the growing oocytes. The resulting MATMUT embryos lack maternal EHMT2 proteins in the postnatal oocyte, zygote, 1-cell, and early two-cell stages, but regain zygotic EHMT2 from the normal paternal allele, inherited from the sperm, during embryo development. Control *Ehmt2*^*fl/+*^ EPCs (MATWT) were obtained by crossing *Ehmt2*^*fl/fl*^ females in the absence of *Zp3-cre* with JF1 males. Because of variable developmental delay of the at the *Ehmt2*^*mat*−/+^ MATMUT embryos we used stage-matched the EPCs from 6-somite embryos.

Maternal *Ehmt2* mutation caused the loss of imprinting of *Gab1*, and *Platr20* in all of the MATMUT EPCs (Fig. 1A). Other genes (*Smoc1*, *Sall1*, *Jade1*, *Platr4*) showed relaxation of imprinting in a few but not all MATMUT EPCs (Fig. 1A). However, *Slc38a4* maintained its paternal allele-specific expression in all MATMUT EPCs. The bias and the expression levels of those genes with loss or relaxation of non-canonical imprinting only slightly changed in the MATMUT EPCs compared to MATWT EPCs (Fig. 1A and B), indicating that the maternal effect of *Ehmt2* is weaker than its zygotic effect. These results revealed that maternal *Ehmt2* mutation selectively and variably affects the paternal allele-specific expression of non-canonical imprinted genes in the EPC.

### The ERVK promoters of non-canonical imprinted genes are suppressed by EHMT2

In the extra-embryonic tissues, the main transcripts of most non-canonical imprinted genes initiate from alternative promoters that coincide with ERVK transposable elements (*10*). The genome browser images of our allele-specific RNA-seq results revealed that the regular promoters of these genes were transcribed at much lower levels than their ERVK promoters. *Gab1* and *Smoc1* are shown as examples in Figure 2. In WT samples, *Gab1* was paternal allele-specifically transcribed from the ERVK promoter, but not from the regular promoter in the EPCs collected from reciprocal mouse crosses (Fig. 2A and Fig. S1). This is in accordance with the paternal allele-specific H3K4me3 signal at the ERVK promoter and biallelic H3K4me3 at the regular promoter in the extraembryonic ectoderm (E×E) at 6.5 dpc (*10*), plotted for comparison. Interestingly, whereas the ERVK-driven *Gab1* and *Smoc1* transcripts were paternally expressed in the WT EPCs they were biallelically expressed in the HOMO EPCs (Fig. 2), and this was true at the 6-somite and 12-somite stages (Fig. S1), revealing that zygotic EHMT2 is required for repressing their RLTR11B/ERVK and RLTR15/ERVK promoters, respectively, in the maternal allele.

**Fig. 2.**
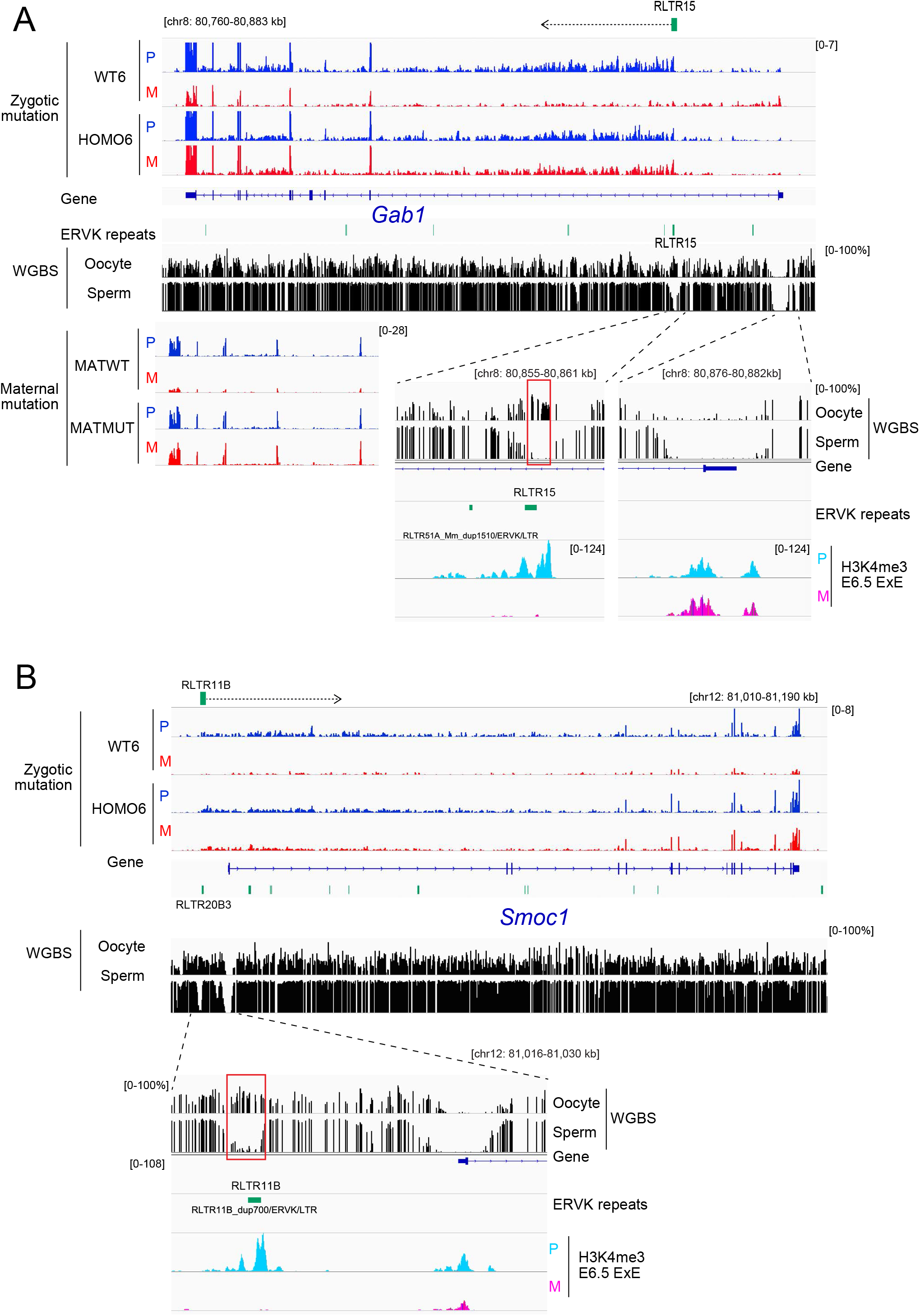
The maternal allele of ERVK-promoters of non-canonical imprinted genes are suppressed by EHMT2 in the ectoplacental cone. **(A)** Derepression of the alternative ERVK-promoter of *Gab1* in the EPC. Genome browser views are shown of the split RNA-seq tracks (RPKM) in the paternal (P) and maternal (M) alleles. One out four *Ehmt2*^−/−^ (HOMO6), and one out of four *Ehmt2*^+/+^ (WT6) EPCs is displayed at the 6-somite stage, representing one of the reciprocal crosses (JF1×129). The WGBS DNA methylation data in the gametes is shown for comparison. The RLTR11B/ERVK promoter-initiated transcription is marked by a dotted arrow. Representative *Ehmt2*^mat−/+^ (MATMUT), and control *Ehmt2*^fl/+^ (MATWT) EPC samples are displayed at the left bottom. The regular and ERVK promoter areas are enlarged at the bottom. Red rectangle marks the germline DMR at the ERVK. The allele-specific H3K4me3 ChIP-seq data in the extraembryonic ectoderm (E×E) (*10*) is shown for comparison. Scales are uniform per replicates and alleles, given in square brackets. **(B)** Derepression of the alternative ERVK-promoter of *Smoc1* in the EPC. Genome browser views are shown as above. The RLTR15/ERVK promoter-initiated transcription is marked by dotted arrow.

We found that similar to *Smoc1* and *Gab1*, the potential novel non-canonical imprinted gene, *Sall1*, has an upstream alternative promoter at an RLTR20B3/ERVK repeat sequence, which carries paternal-specific H3K4me3 in the E×E at 6.5 dpc (Figure 3). Again, the ERVK-driven transcript of *Sall1* was paternally expressed in the WT EPCs but was biallelically expressed in the HOMO EPCs (Fig. 3). In summary, EHMT2 suppressed the maternal allele at the ERVK promoter, but not at the regular promoter at each of these non-canonical imprinted genes. These results indicate that, in the EPC, the alternative ERVK promoters of these non-canonical imprinted genes specifically require zygotic EHMT2 for their repression in the maternal allele.

**Fig. 3.**
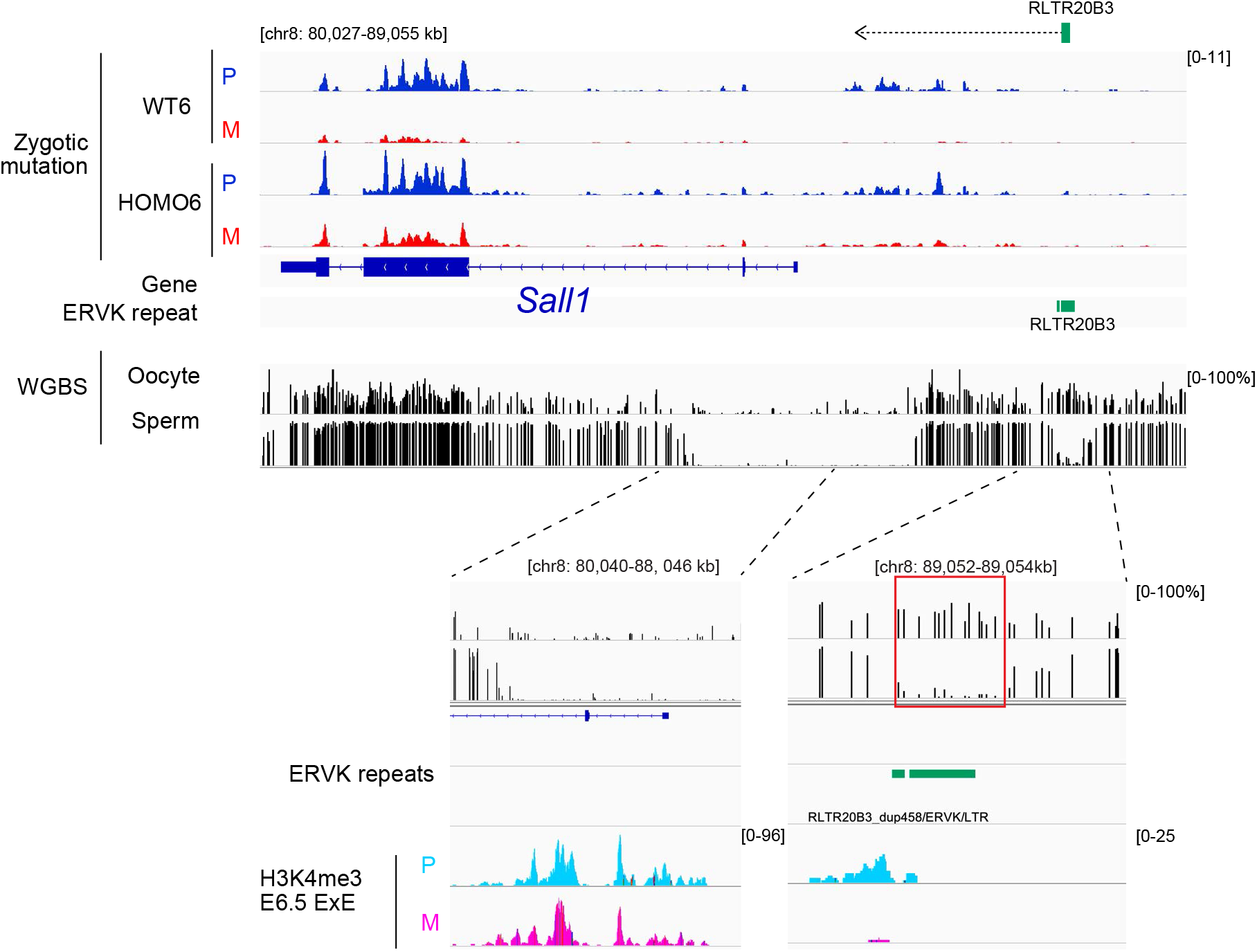
The ERVK promoter of a novel non-canonical imprinted gene, *Sall1* is suppressed by EHMT2 in the paternal allele. Derepression of the alternative ERVK promoter of *Sall1* in the EPC. Genome browser views are shown as explained in Figure 2. The RLTR20B/ERVK promoter-initiated transcription is marked by dotted arrow.

### The ERVK promoters of non-canonical imprinted genes are germ line DMRs

We aligned whole genome bisulfite (WGBS) DNA methylation data from Wang et al. (*23*) to our RNA-seq results, and noticed that the ERVK promoters of *Gab1*, *Smoc1*, and *Sall1* are methylated in the oocyte but are unmethylated in the sperm (Fig. 2 and Fig. 3). In addition, we found that the ERVK promoters at the other non-canonical imprinted genes are also methylated (over 50%) in the oocyte, but unmethylated in the sperm (Fig. S2). This methylation persists into the 2-cell- and the 4-cell stage, but not to the inner cell mass of the blastocyst at 3.5 dpc. These ERVK promoters are, therefore, weak gDMRs, which are not maintained in the preimplantation-stage embryo (Fig. S2).

### Protein-coding non-canonical imprinted genes and ERVK-driven imprinted non-coding RNAs in their proximity synchronously respond to EHMT2

ERVK-driven paternally expressed imprinted non-coding RNAs were detected in the proximity of some of the known non-canonical imprinted genes (*10*). For example, *Gm32885*, driven by an RLTR13F/ERVK repeat is located upstream of *Slc38a4*, and is transcribed in the same orientation as *Slc38a4* (Fig. 4A). Long RNAs *Platr4* and *2400006E01Rik*, which are driven by a RLTR20C/ERVK and RLTR31B/ERVK repeat, respectively, are located upstream of *Jade1 (Pfh17)*, and are transcribed in the opposite direction, away from *Jade1* (Fig. 4B). *Jade1* is transcribed from one of its regular promoters in the EPC, which does not overlap with ERVK elements. We found that, whereas *Gm32885*, *Platr4* and *2400006E01Rik* were expressed exclusively from the paternal allele in the WT EPCs, they became biallelically expressed in the HOMO EPCs (Fig. 1A and Fig. 4A and 4B). These results reveal that EHMT2 synchronously regulates the ERVK-driven imprinted noncoding RNAs with the imprinted genes nearby.

**Fig. 4.**
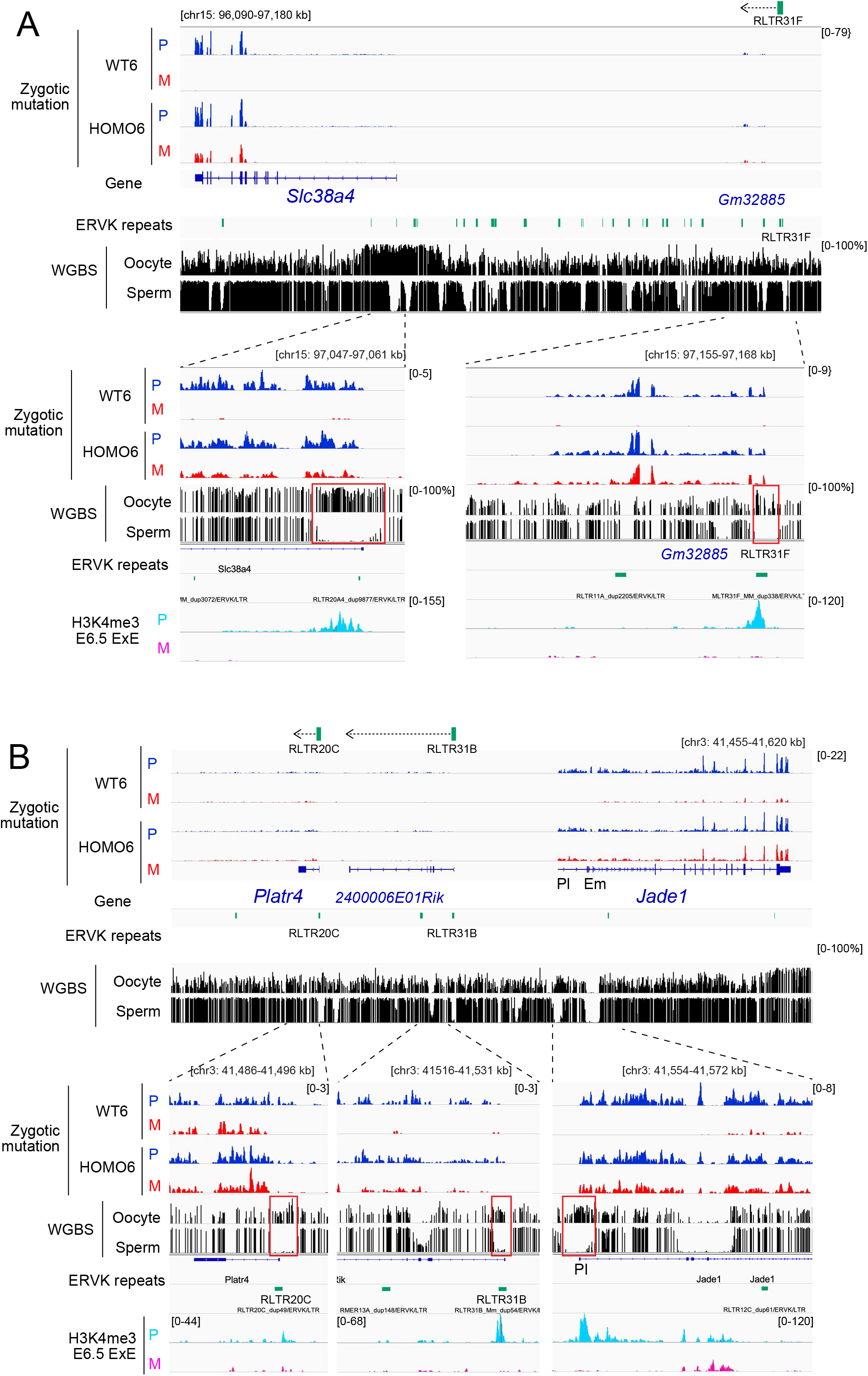
Protein-coding non-canonical imprinted genes and ERVK-driven imprinted non-coding RNAs in their proximity synchronously respond to EHMT2. **(A)** Derepression of the RLTR31F/ERVK promoter-driven noncoding RNA *Gm32885* in the proximity of *Slc38a4* in the EPC. Genome browser views are shown as explained in Figure 2. The RLTR31F/ERVK promoter-initiated transcription is marked by dotted arrow. **(B)** Derepression of the RLTR31B/ERVK promoter-driven noncoding RNA *2400006E01Rik* in the proximity of *Jade1* in the EPC. The location of the RLTR20C/ERVK promoter-driven noncoding RNA, *Platr4*, is also shown. The imprinting of this transcript is variable. Genome browser views are shown as explained in Figure 2. The ERVK promoter-initiated transcripts are marked by dotted arrows. EPC- and embryo-specific regular promoters are marked (Pl, and Em, respectively).

### *Sfmbt2* and its large miRNA gene cluster are regulated by EHMT2 in the EPC

We found that the protein-coding *Sfmbt2* gene, and the upstream located antisense, noncoding RNA gene *Gm13261*, were both paternally expressed in the WT EPCs, but became biallelically expressed in HOMO EPCs (Fig. 5A). In addition, the large miRNA gene cluster inside the intron 10 of *Sfmbt2* showed the same pattern, the miRNA precursor sequences were paternally expressed in WT EPCs as expected (*24*) but became biallelically expressed in HOMO EPCs (Fig. 5A). *Sfmbt2*, Gm13261, and the miRNA gene cluster maintained their paternal allele-specific expression in the MATMUT EPCs (Fig. 5A). The promoter of *Gm13261* overlaps with an RLTR11B/ERVK repeat element, which is a gDMR (Fig. 5B). We also noticed a large number (n=125) of ERVK repeats in the intron 10 of *Sfmbt2*, interspersed with the miRNA genes (Fig. 5B). These results collectively reveal that EHMT2 is required for the maintenance of paternal allele-specific expression in this imprinted gene domain in the EPC. Suppression of the ERVKs (RLTR11B/ERVK or in the miRNA cluster) by EHMT2 may be required for this process.

**Fig. 5.**
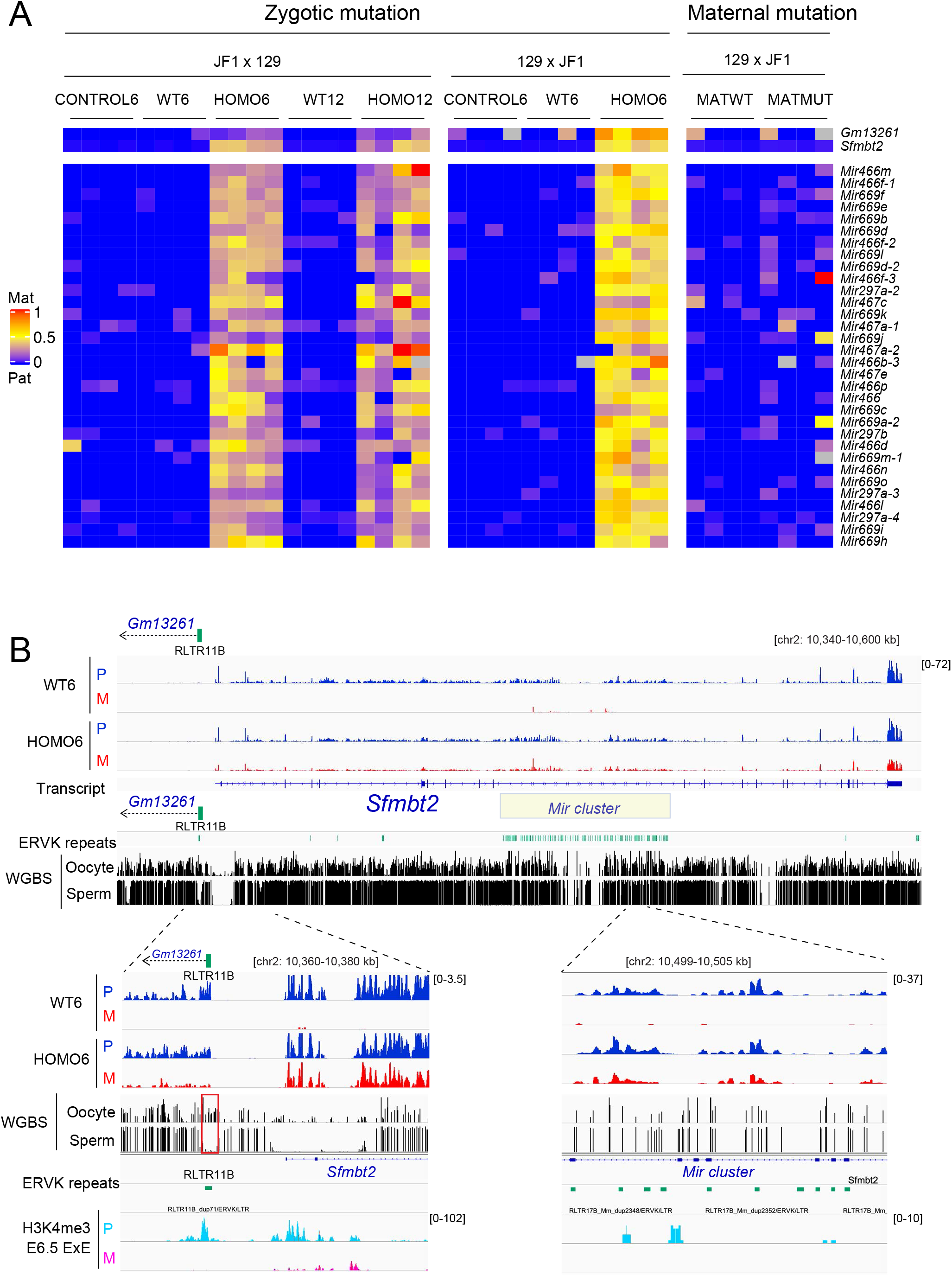
*Sfmbt2* and its large miRNA gene cluster are regulated by EHMT2 in the EPC. **(A)** Allelic expression ratios (Mat/Mat+Pat) of the *Sfmbt2* gene and its intronic miRNAs across all EPC samples. *Gm13261* is an ERVK-driven antisense non-coding RNA located upstream of the *Sfmbt2* gene promoter. Other details are described in Fig. 1A. **(B)** Genome browser views of the *Sfmbt2* locus. The details are the same as described in Figure 2. The large miRNA cluster is marked with a rectangle. Note that it corresponds to a large ERVK cluster. The promoter area and a detail from the miRNA cluster are enlarged at the bottom.

### EHMT2 is required for biallelically repressing the ERVK promoters of non-canonical imprinted genes in the embryo

Noncanonical imprinted genes are biallelically expressed from their regular promoters in the 6.5 dpc epiblast and later in the embryo, and their ERVK promoters are biallelically suppressed by somatically established DNA methylation (*10*). We also tested whether EHMT2 plays a role in regulating noncanonical imprinted genes in the embryo. In the WT embryos the alternative ERVK promoters of *Gab1*, *Smoc1* and *Sall1* were silenced in both maternal and paternal alleles, while their regular promoters were active in both maternal and paternal alleles, as displayed for *Smoc1* in Figure 6A. Interestingly, in HOMO embryos the ERVK promoters of these genes were aberrantly derepressed in both maternal and paternal alleles. According to our WGBS results in 9.5 dpc embryos, biallelic derepression in the HOMO embryos coincided with a loss of DNA methylation at the corresponding ERVKs. The fully methylated RLTR11B/ERVK promoter upstream of *Smoc1* in WT embryos changed to a fully unmethylated state in HOMO embryos (Fig.6A), which is consistent with losing methylation in both parental alleles.

**Fig. 6.**
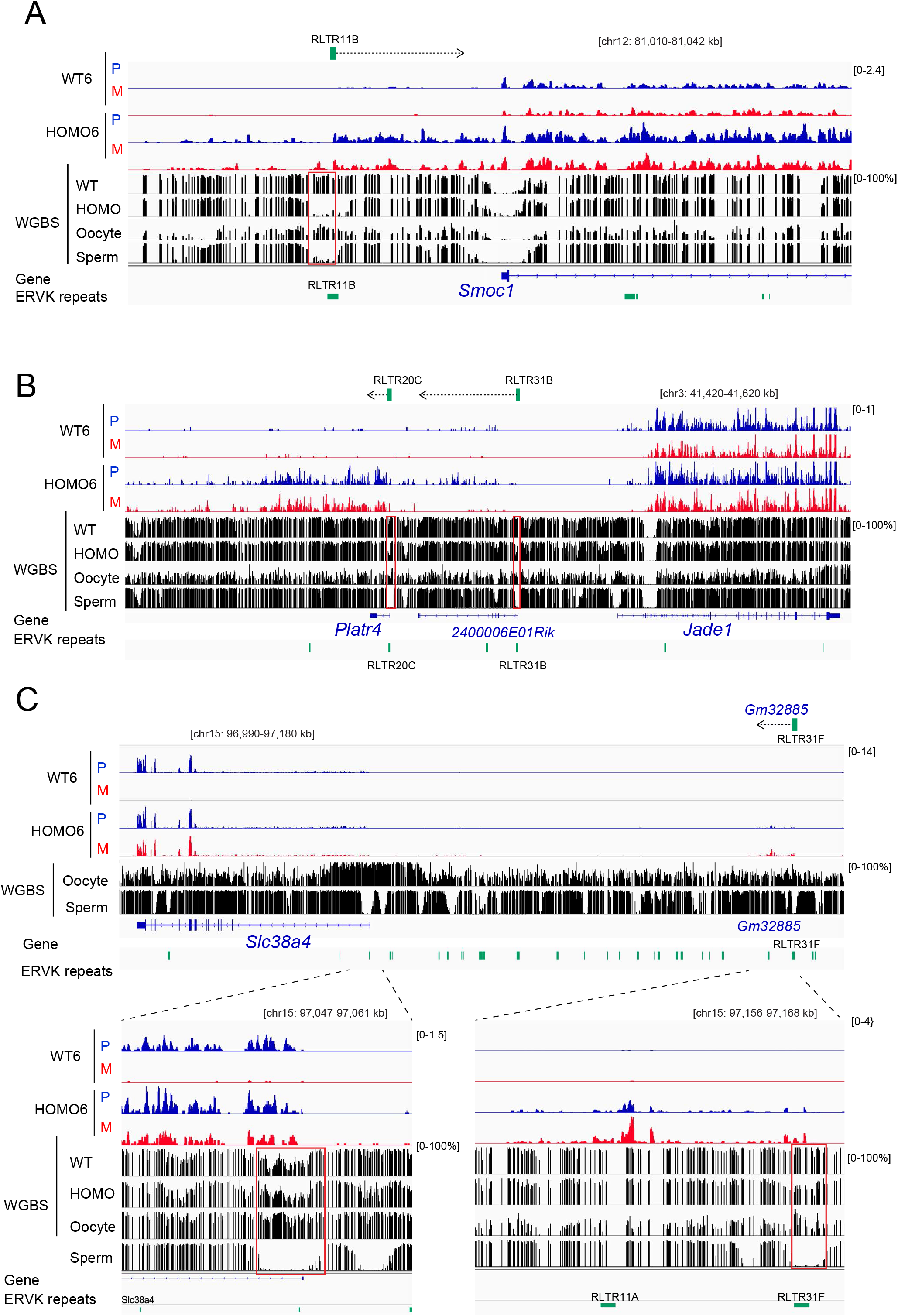
EHMT2 is required for biallelically repressing the ERVK promoters of non-canonical imprinted genes in the embryo. **(A)** Biallelic activation of the ERVK promoter of *Smoc1* in HOMO6 embryos. Genome browser views are shown as described in Figure 2, except WGBS methylation results are added from WT and HOMO 9.5 dpc embryos from our own experiments. Note the loss of CpG methylation at the RLTR11B/ERVK promoter. **(B)** The RLTR20C/ERVK and RLTR31B/ERVK promoters driving *Platr4* and *2400006E01Rik,* respectively lose DNA methylation and are biallelically derepressed in HOMO6 embryos. **(C)** Derepression of the maternal allele of *Slc38a4* gene in HOMO6 embryos coincides with the biallelic derepression of the upstream RLTR31F/ERVK-driven noncoding RNA *Gm32885*. The promoter areas are enlarged at the bottom.

The two ERVK-driven imprinted noncoding RNA genes *Platr4* and *2400006E01Rik* located upstream of *Jade1* were silent in WT embryos but were derepressed in HOMO embryos, and this corresponded to decreased DNA methylation at the RLTR20C/ERVK and RLTR31B/ERVK promoters, respectively (Fig. 6B). The *Jade1* gene was biallelically expressed from the embryo-specific regular promoter, while its EPC-specific promoter was biallelically silenced in WT embryos. Transcription out of the *Jade1* regular promoters did not change in the HOMO embryos.

Among non-canonical imprinted genes that are paternally expressed in the extraembryonic tissues, and do not depend on gametic DNA methylation, there is one exception, *Slc38a4*, which is paternally expressed in the embryo as well, and does depend on a gDMR at the *Slc38a4* promoter, which is established in the oocyte (*25*). We found that *Slc38a4* expression became biallelic in HOMO embryos (Fig. 6C), consistent with another study, which showed the upregulation of the *Slc38a4* transcript in *Ehmt2*^−/−^ embryos (*15*). However, the majority of CpGs in the *Slc38a4* gDMR maintained their high DNA methylation level between the WT and HOMO embryos. The upstream RLTR31F/ERVK-driven imprinted RNA gene, *Gm32885*, was biallelically derepressed in the maternal allele in HOMO embryos and DNA methylation was reduced at this ERVK (Fig. 6C).

The *Ehmt2* mutation eliminated the catalytic SET domain of EHMT2 (*22*). The simplest explanation for EHMT2-s effect at the non-canonical imprinted genes is that H3K9me2 levels were insufficient in the mutant embryos at the ERVKs at non-canonical imprinted genes. This implies that WT embryos have H3K9me2 at those sequences. We examined these regions in publicly available ChIP-seq datasets (*15, 26, 27*) and show genome browser images in Figure S3. We found that H3K9me2 shows enrichment at the ERVKs at the non-canonical imprinted genes in the body and head of mouse embryos (*26*) and in epiblast stem cells (*27*), which correspond to the 6.5 dpc epiblast. This enrichment is consistent with our finding that these ERVK-s require EHMT2 catalytic activity for their suppression in the embryo.

By examining a published dataset (*2*) we found that ES cells that carry mutation of *Ehmt2* display DNA hypomethylation at those regions, together with an increase in the enrichment for the H3K4me3 mark (Fig. S4). This is consistent with the possibility that the effect EHMT2 on ERVK DMR methylation may take place as early as the blastocyst stage.

These results collectively show that EHMT2 suppresses both alleles of the alternative ERVK promoters of non-canonical imprinted genes in the embryo, and this suppression requires the histone methyltransferase activity of EHMT2 and also involves DNA methylation at the ERVKs. The same is true for the ERVKs that are synchronously regulated in the proximity of non-canonical imprinted genes.

### EHMT2 has a minor contribution to maintaining canonical imprinting

It was previously reported that the imprinting of a few maternally expressed genes along distal chromosome 7 was lost in the absence of EHMT2 in the placenta, but not in the embryo, and that gDMR DNA methylation was not affected (*21*). Our dataset revealed that some maternally expressed imprinted genes along distal chromosome 7, such as *H19*, *Ascl2*, *Tssc4*, *Trpm5*, *Slc22a18*, *Phlda2*, and *Cdkn1c,* showed significantly altered allelic expression ratios (FDR<0.05) in HOMO versus WT EPCs (Fig. S5 and Table S2). The relaxation of the paternal allele was very slight for *H19* and *Cdkn1c*, while it was more robust for others, such as *Ascl1* and *Phlda2* (Fig. S5B). Maternal *Ehmt2* mutation did not affect these maternally expressed genes (Fig. S5A). These data confirmed that zygotic EHMT2 contributes to the repression of the paternal alleles of maternally expressed genes on distal chromosome 7 and identified additional genes compared to those reported earlier (*21*).

## Discussion

With the exception of *Slc38a4*, non-canonical imprinted genes lack domain-wide regulation by an imprinting control region (ICR). The imprinting of these genes in the extraembryonic tissues may depend on their alternative ERVK promoters, or the synchronously imprinted ERVK promoters in their vicinity. Our finding, that EHMT2 is essential for proper suppression of the alternative ERVK promoters and synchronously imprinted ERVK transcripts at non-canonical imprinted genes reveals a novel essential mechanism for their parental allele-specific suppression beyond preimplantation, when the H3K27me3 mark is lost. EHMT2 was required to suppress each of the non-canonical imprinted genes maternal allele-specifically in the EPC and biallelically in the embryo.

Using zygotic mutation of *Ehmt2* we found that EHMT2 is required for maintaining paternal allele-specific expression of non-canonical imprinted genes. It is perhaps the easiest to envision how an alternative ERVK promoter can regulate the host promoter in cis. Indeed, when the ERVK promoter of *Gab1* was deleted in the paternal allele, the allelic expression bias of *Gab1* was reduced (*10*). However, the oppositely transcribed upstream ERVK promoters or ERVK promoters at a distance may require different mechanisms. They may rely on the shared chromatin environment in a domain. Alternatively, long-range interactions between the active paternal allele of the ERVK and the regular promoter may be at play, and EHMT2 provides suppression for such interaction in the maternal allele. It will be interesting to test whether these ERVK elements and/or the ERVK-driven imprinted noncoding RNAs, such as *Gm32885* located upstream of *Slc38a4*, *Platr4* and *2400006E01Rik* located upstream of *Jade1*, and *Gm13261* located upstream *Sfmbt2* can play a similar role in maintaining the activity in the paternal allele at their respective protein-coding non-canonical imprinted gene.

By generating maternal mutants, we found that maternal EHMT2 in the oocyte/zygote is essential for the paternal-allele-specific expression of some non-canonical imprinted genes in the EPC, and this maternal effect cannot be corrected by the zygotic expression of *Ehmt2*. Maternal *Ehmt2* mutation caused the loss of imprinting of *Gab1* and the relaxation of imprinting of *Smoc1* and *Sall1* (Fig. 1), indicating that their ERVK promoters may have already been programmed by maternal EHMT2 in the oocytes, or that EHMT2 was required for maintaining the suppressive mark in the zygote/early embryo. This is consistent with our earlier study, where we showed by immunocytochemistry that EHMT2 is required in the zygote’s maternal pronucleus for resisting TET-mediated oxidation (*22*). It will be interesting to test whether removing the ERVKs at non-canonical imprinted genes in the growing oocyte will affect the allele-specific expression of these genes in the extraembryonic tissues.

The suppressing effect of EHMT2 at canonical imprinted genes is subtle compared to what is observed at non-canonical imprinted genes. This is likely because canonical imprinted genes can rely on the continuity of gDMR-dependent mechanisms of suppression, while non-canonical imprinted genes depend on a cascade of events and on DMRs where DNA methylation is not continuously maintained from gamete to soma.

The paternal allele-specific expression of non-canonical imprinted genes depends on a combination of sequential events. The primary mark, broad H3K27me3, is gradually lost at non-canonical imprinted genes during preimplantation development (*11*). DNA methylation at the ERVKs is also lost by the 4-cell stage (Fig. S2). Our study revealed that the catalytic activity of EHMT2 is an additional layer of regulation and H3K9me2 is likely part of this picture. Zygotic *Ehmt2* expression and genome-wide chromatin enrichment in H3K9me2 may increase during the pre- to post-implantation transition and H3K9me2 may take over the maternal H3K27me3 signal and suppress the maternal allele directly. Paternal-allele-specific H3K4me3 (*10*) may antagonize H3K9me2 deposition in the extraembryonic tissues, but not in the embryo, hence the biallelic suppression of ERVKs in the embryo. We show that EHMT2-dependent biallelic suppression of the ERVKs in the embryo involves DNA methylation. It is likely that EHMT2-dependent suppression of ERVKs in the EPC similarly involves DNA methylation, but only in the maternal allele. Future studies including ChIP-seq are needed to find out whether, and how H3K9me2 marking targets DNA methylation to the ERVK promoters.

Placenta development is affected in *Ehmt2* mutant mice (*21*). Our results suggest that EHMT2’s broad effect on placenta development may involve the regulation of non-canonical and canonical imprinted genes. Further studies are needed to find out the role of imprinting at specific non-canonical imprinted genes in regulating placentation and fetal growth. A targeted deletion of a 53 kb region encompassing the entire miRNA cluster in the tenth intron of *Sfmbt2* caused placental defects and partial embryonic lethality (*24*), but is unknown how loss of imprinting, and the resulting biallelic overexpression of this cluster affects development of the placenta and the embryo. Similarly, the absence of *Gab1* is deleterious to placenta development (*28, 29*), but the effect of its overexpression is not known.

The presence of a long miRNA cluster in the intron 10 of *Sfmbt2* correlates with its imprinting and it was hypothesized to drive imprinting at this locus (*30*). Another explanation is equally possible, that the ERVK repeats embedded in the miRNA cluster drive its imprinting, and EHMT2 may be involved in this process. On a similar note, species-specific repeats have been shown to underlie the evolution of imprinted gDMRs (*25*), for example an MT2 repeat is required for setting up the maternal gDMR at the *Slc38a4* gene in the mouse. The gDMRs at the ERVKs may also be involved in the evolution of non-canonical imprinting. This would imply that gDMRs at the ERVK promoters are required for non-canonical imprinting, which is inconsistent with the findings using maternal knockouts of *Dnmt3a/3b* (*10*) and *Dnmt3l* (*11*). However, one germline-specific de novo DNA methyltransferase has not been accounted for. DNMT3C specifically methylates repeat elements, including LINEs and ERVKs in the male germline (*31, 32*), and may also play such a role in the female germline. Even though the ovary has low level of *Dnmt3c* transcription, this may be different for growing oocytes. ERVKs may be expressed in the male germline but not in the female germline, and the corresponding H3K4me3 marking at those active promoters may repel de novo DNA methyltransferases in prospermatogonia but not in growing oocytes. This is consistent with the finding that DNA methylation occurred by default in prospermatogonia except where antagonized by H3K4 methylation (*33*). H3K27me3, on the other hand would suppress the ERVKs in the growing oocytes, hence H3K4me3 would not antagonize DNA methylation. Marking ERVKs in the oocyte by DNA-methylation would provide a narrow, locus-specific non-canonical imprint in addition/or as opposed to the very broad H3K27me3 mark.

In summary, we provide evidence that EHMT2 catalytic activity is a novel essential component of the mechanism that regulates non-canonical imprinted genes in the extraembryonic tissues and in the embryo.

## Materials and Methods

All animal experiments were performed in accordance with the National Institutes of Health Guide for the Care and Use of Laboratory animals, and the Institutional Care and Use Committee-approved protocols at Van Andel Institute (VAI).

### Mouse models

An *Ehmt2*^*fl/fl*^ conditional knockout mouse line was generated in our laboratory by gene targeting in 129S1/SvImJ ES cells (*22*). The floxed SET domain of *Ehmt2* was removed to generate the *Ehmt2*^*+/−(129S1)*^ heterozygous mouse line by crossing the *Ehmt2*^*fl/+*^ male with 129S1/Sv-Hprt^tm1(CAGcre)Mann/J^ transgenic female mice. *Ehmt2*^*fl/fl*^ and *Ehmt2*^*+/−(129S1)*^ mice were maintained in the 129S1 genetic background. The *Ehmt2*^*+/−(129S1)*^ mice were crossed with wide-type JF1/Ms (JF1) mice for more than ten generations to obtain the *Ehmt2*^*+/−(JF1)*^ heterozygous mouse line, in which most of the genome has been replaced by JF1 chromosomes during meiotic recombination except that the *Ehmt2* locus and its close neighbors in the approximate interval of chr17:34647146-35241746. The *Ehmt2*^*+/−(JF1)*^ heterozygous mouse line was maintained in the JF1 genetic background.

### Collection of *Ehmt2*^*−/−*^ and *Ehmt2*^*mat−/+*^ EPCs, embryos, and their controls

Reciprocal crosses were set up between *Ehmt2*^*+/−(129S1)*^ and *Ehmt2*^*+/−(JF1.N10)*^ heterozygous mice to obtain embryos by natural mating. Crossing *Ehmt2*^+/−(JF1.N10)^ females with *Ehmt2^+/^*^−(129S1)^ males yielded *Ehmt2*^−(JF1)/−(129S1)^ zygotic mutant and *Ehmt2*^+(JF1)/+(129S1)^ wild type F1 embryos and EPCs. The reciprocal cross resulted in *Ehmt2*^−(129S1)/−(JF1)^ mutant and *Ehmt2*^+(129S1)/+(JF1)^ wild type samples. Control *Ehmt2*^+(129S1)/+(JF1)^ samples were obtained by crossing wild type 129S1 and JF1 male mice. Control *Ehmt2*^+(JF1)/+ (129S1)^ samples were obtained by crossing wild type JF1 female and 129S1 male mice. Natural matings were set up between *Ehmt2*^fl/fl^; *Zp3-cre*^*Tg/0*^ females and wild type JF1 males to generate *Ehmt2*^*mat−/+JF1*^ maternal mutant samples. Their control samples came from crossing *Ehmt2*^*fl/fl*^; *Zp3-Cre*^*Tg-*^ females with wild type JF1 males. The EPCs and embryos were dissected at different time points, on 8 dpc at 9 am, 8 dpc at 6 pm and 9 dpc at 9 am as summarized in Table S1, and they were flash-frozen on dry ice in an aliquot of 100 μL of RNA-Bee (Tel-Test). Genotyping for *Ehmt2* mutation status and sex was done by PCR from the allantois as described earlier (*22*).

### RNA isolation and RNA sequencing

Total RNA was isolated from individual EPCs using RNA-Bee (Tel-Test) and chloroform extraction followed by isopropanol precipitation. Genomic DNA was removed by treating the total RNA samples with the DNA-free Kit (Ambion) followed by ethanol precipitation. The quality and integrity of the total RNA samples were checked on the Fragment Analyzer (Advanced Analytical Technologies, Inc.). 500 ng total RNA was used for generating strand-specific RNA-seq libraries using the KAPA Stranded RNA-Seq Kit with RiboErase (Kapa Biosystems, MA). RNA-seq libraries were sequenced on an Illumina NextSeq 500 platform with paired-end 75 bp read length.

### RNA-seq data analysis

Reads were aligned to the mm10 reference genome using STAR (version 2.6.0c). Using the Ensembl gene annotation (v82), counts per gene was calculated using featureCounts. Genes with CPM > 1 in at least two of the samples were retained for further analysis. Differential gene expression was performed using edgeR (*34, 35*).

### Imprinting analysis

The SNP information between the 129S1 and JF1/Ms mouse strains were extracted using a custom python script from the mouse genomic variation data downloaded from the Sanger Institute (https://www.sanger.ac.uk/data/mouse-genomes-project/). To minimize the alignment bias, all SNPs between the 129S1 and JF1 strains were masked as “N” to generate a custom SNP-masked mm10 genome. Paired-end reads were aligned to this custom genome using Hisat2 with parameters “–sp 1000,1000 –dta-cutfflinks”. Parental allele-specific reads were extracted using SNPsplit. Gene counts per Ensembl transcript on the maternal or paternal alleles was calculated using featureCounts. Allelic expression ratios were calculated using edgeR for all genes with CPM > 1 in at least two of the samples. Significant changes in imprinting was determined using cutoff value of FDR < 0.05.

### Whole genome bisulfite sequencing (WGBS) analysis

WGBS analysis was used to map DNA methylation in embryo DNA at 9.5 dpc. One sample contains DNA from two *Ehmt2* homozygous mutant embryos (one male and one female combined), and the control sample contains DNA from a wild type female embryo. In order to generate the whole genome libraries with bisulfite converted DNA, embryo DNA was sonicated to approx. 150 bp DNA fragments. Further, DNA was end repaired by using End-It- DNA End-Repair Kit (Epicentre) and linker ligated with T4 ligase (NEB). The linked ligated DNA was bisulfite converted using EZ DNA Methylation-Gold™ Kit according to manufacturer’s instructions (Zymo Research) and amplified with Pfu Turbo polymerase (Agilent). Sequencing was performed in the Illumina HiSeq 2500 platform at the Integrative Genomic Core at the City of Hope Cancer Center. Pair-end reads were aligned to the mm10 genome using Biscuit https://huishenlab.github.io/biscuit/) and duplicates were marked using Picard’s MarkDuplicates. DNA methylation and genetic information were extracted using biscuit’s pileup and CpG beta values were extracted using vcf2bed.

### Statistical analysis and data visualization

Statistical analyses were performed in R (https://www.r-project.org/). The heatmaps in this study were generated using the R package “ComplexHeatmap”. The gene expression data in Fig. 1B was plotted using the R package “ggplot2”. The sequencing tracks were visualized using the Integrative Genomics Viewer (http://software.broadinstitute.org/software/igv/).

## Acknowledgments

We are grateful to Drs. Gavin Kelsey and Courtney Hanna for sharing the H3K4m3 allele-specific analysis. We thank Gerd Pfeifer and Darrell Chandler for their comments on the manuscript. We thank the VAI Genomics Core for RNA deep sequencing of the RNA samples and the VAI Vivarium for mouse maintenance and husbandry. The technical assistance of Yanli Su is greatly appreciated. We acknowledge the funding from VAI.

## Author contributions

Conceptualization (PES); Formal analysis (NP); Funding acquisition (PES); Investigation (TZ, NP, PES); Methodology (PES); Supervision (PES); Validation (TZ, NP, PES); Visualization (TZ, NP, PES); Writing-original draft (TZ); Writing-review and editing (TZ, NP, PES).

## Data availability

The RNA-seq datasets generated in this study have been deposited in the Gene Expression Omnibus under accession number GSE160455 and GSE156538. Allele-specific H3K4me3 ChIP-seq data in E6.5 E×E are based on GEO submission GSE124216. WGBS data of sperm and oocyte were obtained from GSE56697.

## Supplementary Materials

**Fig. S1.**
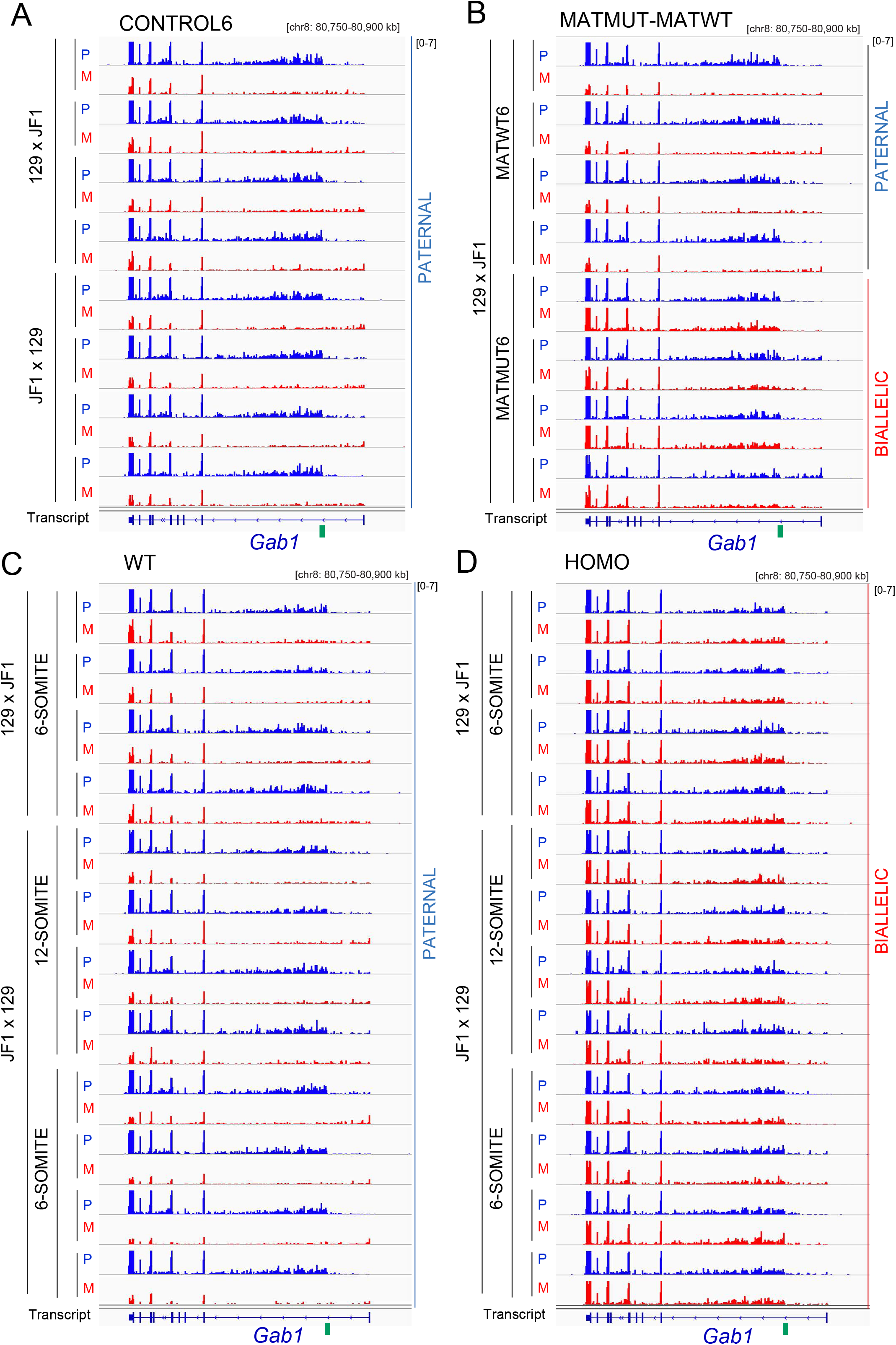
Reproducibility of the RNA-seq results in replicate EPC samples. **(A)** CONTROL EPC samples are displayed in four 6-somite stage EPCs each in two reciprocal crosses (129×JF1 and JF1×129). Split-alleles are shown of *Gab1* as described in Figure 2. Green box shows the location of the ERVK promoter. The scale is set to visualize the derepression from the normally silent maternal (M) allele initiated from the ERVK promoter, while the active paternal (P) allele is saturated. It is fully visible at the scale of [0-27], as shown in the bottom panel in Figure 2A. The samples are described in Table S1. **(B)** Maternal mutant (MATMUT) and maternal wild type (MATWT) EPC sample replicates are shown in the 129×JF1 cross. **(C)** Wild Type (WT) EPC samples. Four 6-somite stage 129×JF1, four 12-somite stage JF1×129, and four 6-somite-stage JF1×129 samples are shown. **(D)** Homozygous zygotic mutant (HOMO) EPC samples. Four 6-somite stage 129×JF1, 12-somite stage JF1×129, and four 6-somite-stage JF1×129 samples are shown. Note the change from paternal to biallelic expression in the mutant samples.

**Fig. S2.**
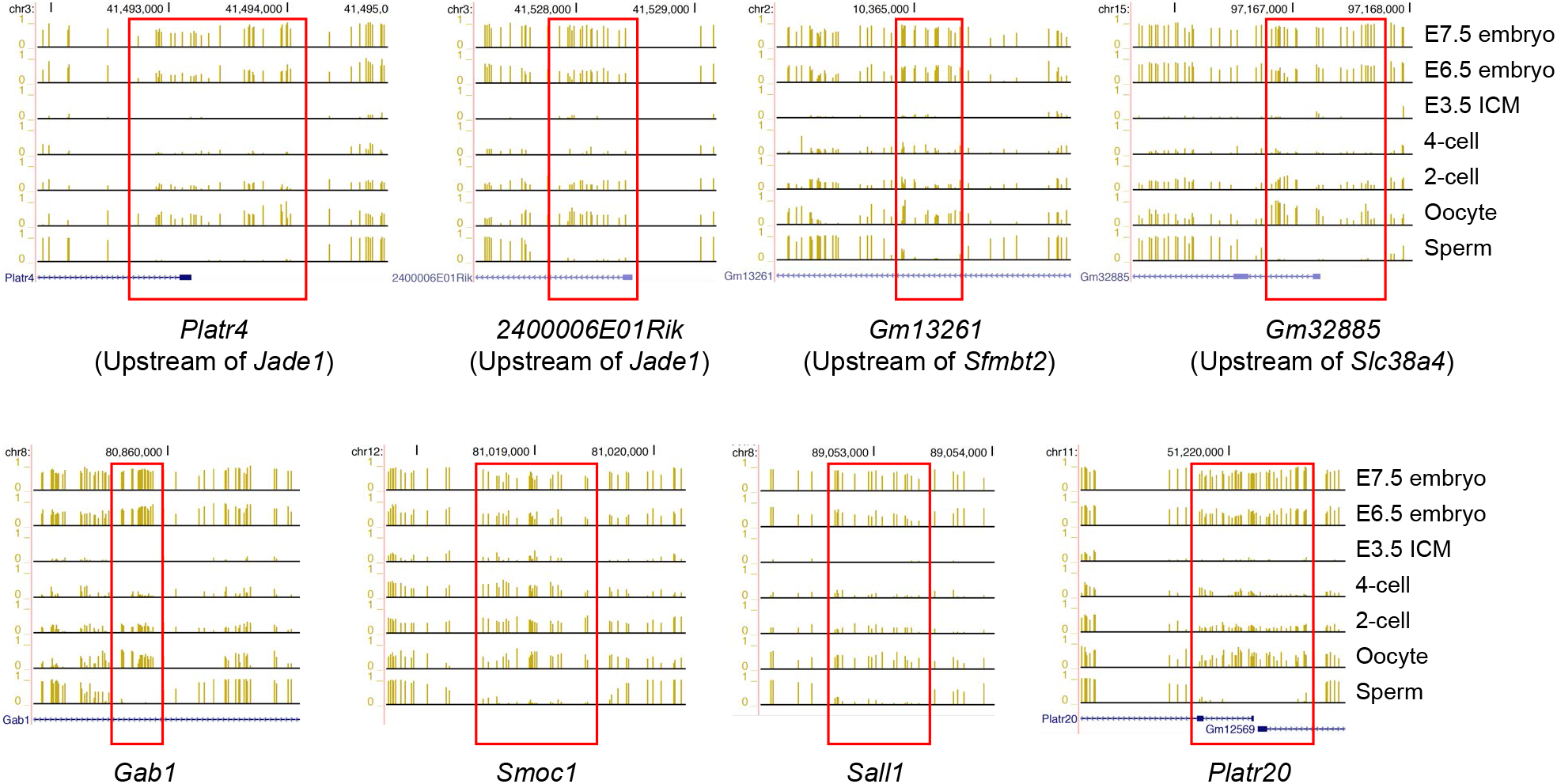
The ERVK promoters of non-canonical imprinted genes are germline DMRs. Genome browser views of DNA methylation data (*23*) are displayed at the ERVK promoters, associated with non-canonical imprinted genes in the oocyte, sperm, 2-cell and 4-cell embryos, E3.5 ICM, E6.5 and E7.5 embryos. Oocyte versus sperm methylation difference at gametic DMRs (rectangles) is not maintained during preimplantation development.

**Fig. S3.**
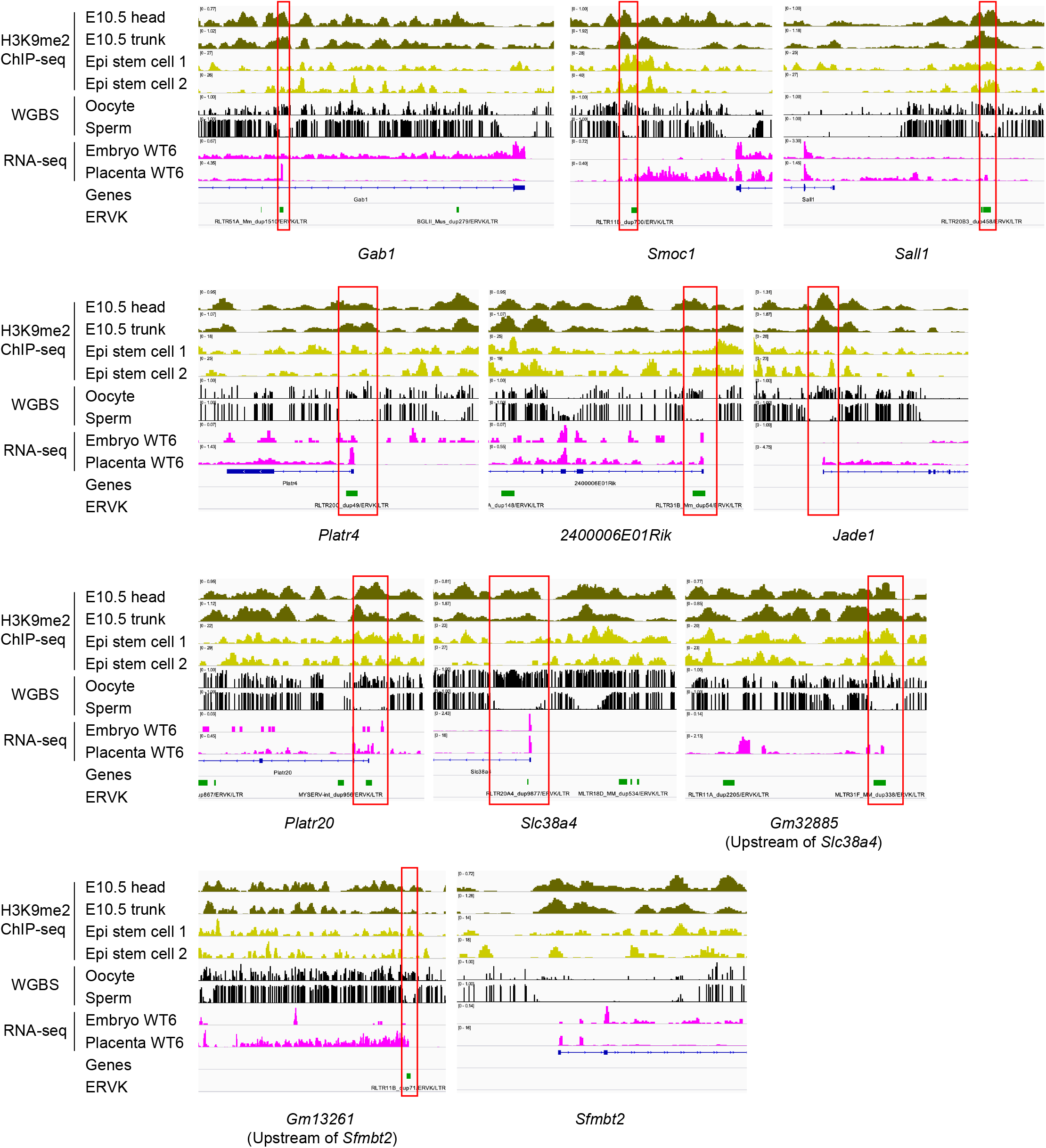
Non-canonical imprinted genes display H3K9m2-rich chromatin at their ERVK promoters. Genome browser views of H3K9me2 ChIP-seq data in epiblast (Epi) stem cells (*27*) and in the head, and the body of 10.5 dpc embryo (*26*) at the ERVK repeats associated with non-canonical imprinted genes. These regions have H3K9me2 enrichment which can respond to inactivating EHMT2 catalytic activity in HOMO embryos. Genome browser views of DNA methylation data (*23*) is also displayed in the oocyte, and sperm together with our RNA-seq results in WT6 embryo and EPC. Note also the different transcription start sites and transcription levels (scales in square brackets) in the EPC and embryo.

**Fig. S4.**
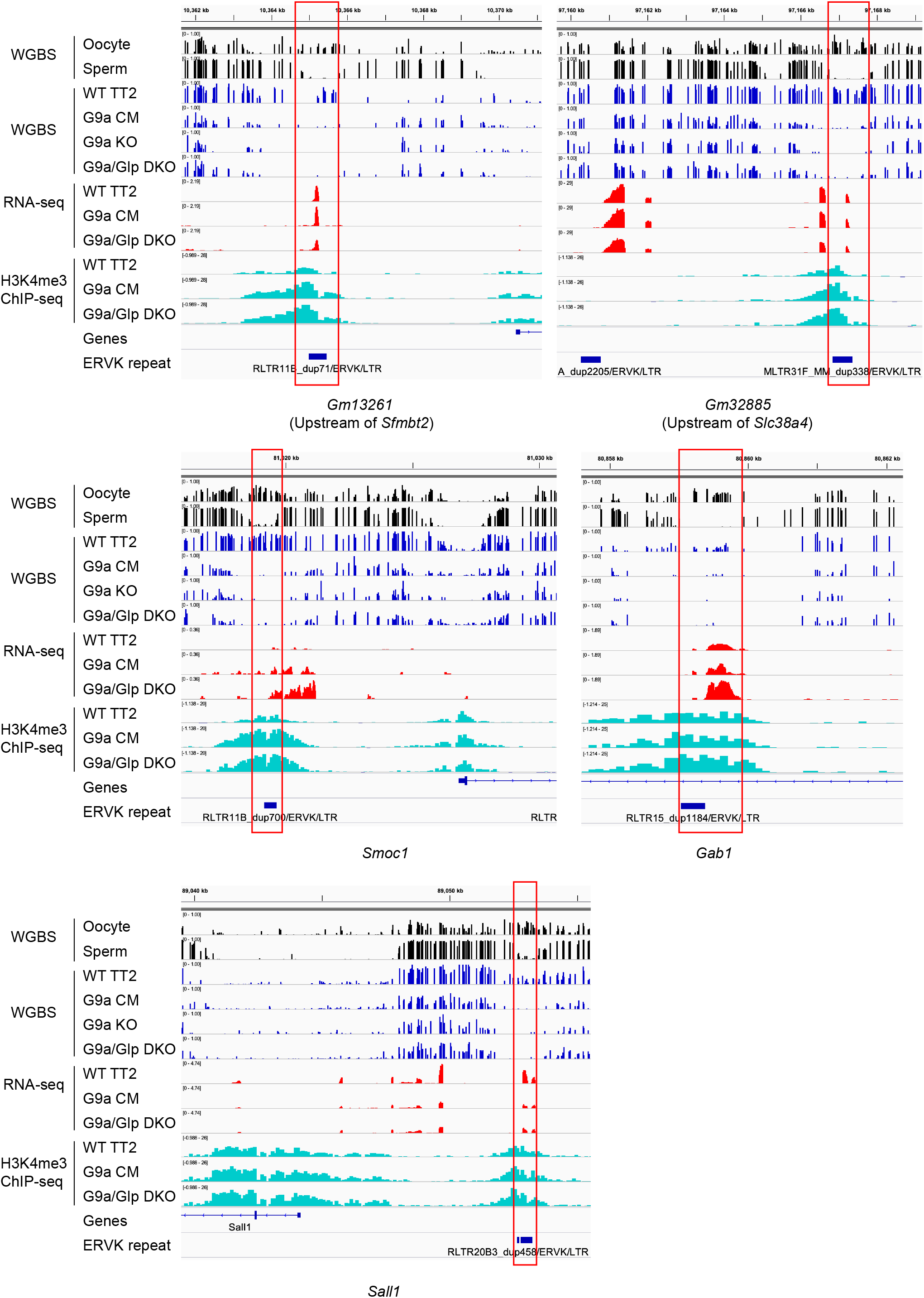
EHMT2 affects chromatin at the ERVK germline DMRs in ES cells. Genome browser views display WGBS DNA methylation data (*23*) at the ERVK promoter gametic DMRs promoters of non-canonical imprinted genes in the oocyte and sperm, Oocyte versus sperm methylation difference (gametic DMR) is marked with rectangles. The other browser lanes show WGBS, RNA-seq, and H3K4me3 ChIP-seq results (*2*) in ES cells of four genotypes: wild type (TT2), *Ehmt2*/*G9a* catalytic mutant (G9a CM), *G9a* knockout (G9a KO) and *G9a*/*Glp* double knockout (DKO).

**Fig. S5.** EHMT2 has a minor contribution to maintaining canonical imprinting in the EPC. **(A)** Allelic expression ratio (Mat/Mat+Pat) of selected maternally expressed canonical imprinted genes located along distal chromosome 7 in the EPC samples as described in Fig. 1A. **(B)** Slight derepression of the paternal allele of canonical imprinted genes. Genome browser images are shown as described in Figure 2.

**Table. S1. List of all EPC samples used in this study.**

**Table. S2. Allelic expression ratios of all imprinted genes in the EPC samples.**

